# Modular organization of synapses within a neuromere for distinct axial locomotion in *Drosophila* larvae

**DOI:** 10.1101/2024.06.29.601329

**Authors:** Kazushi Fukumasu, Akinao Nose, Hiroshi Kohsaka

## Abstract

The ability to generate diverse patterns of behavior is advantageous for animal survival. However, it is still unclear how interneurons in a single nervous system are organized to exhibit distinct motions by coordinating the same set of motor neurons. In this study, we analyze the populational dynamics of synaptic activity when fly larvae exhibit two distinct fictive locomotion, forward and backward waves. Based on neurotransmitter phenotypes, the hemi-neuromere is demarcated into ten domains. Calcium imaging analysis shows that one pair of the domains exhibits a consistent recruitment order in synaptic activity in forward and backward waves, while most other domains show the opposite orders in the distinct fictive locomotion. Connectomics-based mapping indicates that these two domains contain pre- and post-synaptic terminals of interneurons involved in motor control. These results suggest that the identified domains serve as a convergence region of forward and backward crawling programs.

## Introduction

Generating multiple patterns of behavior is critical for animal survival. How multiple motor programs are implemented in a single central nervous system (CNS) is a fundamental problem in behavioral neuroscience. Both dedicated and multi-functional neurons have been identified to be involved in distinct motor functions (1–8). However, how dedicated and multi-functional motor circuits are organized remains still unclear.

Drosophila larval locomotion provides an ideal model system to study multi-functional motor circuits (9–12). Fly larvae show forward crawling by sequential contraction of the segments from posterior to anterior. By propagating the contraction in the opposite direction, larvae crawl backward. Connectomics research and genetic analyses have identified key interneurons that regulate forward and backward crawling (10,12). However, the mesoscopic picture of how dedicated and multi-functional interneurons for forward and backward locomotion are deployed in the CNS is still unclear.

In this study, we identify intrasegmental synapse structures in the CNS. First, we develop a template coordinate to map synapse data into a standardized framework. Then, we analyze the distribution of neurotransmitters in the VNC and find two types of geometric structures: stripes and blobs. We notice that the neuromere can be demarcated into 10 domains. To understand the functional significance of the domains, we conducted calcium imaging of the nervous system. We find one pair of the domains exhibits consistent recruitment order between forward and backward fictive locomotion while the other pairs do not. Connectomics-based analysis reveals interneurons involved in the two domains, PI and AIc, and these neurons include premotor neurons reported previously to be involved in motor control. These results indicate the PI domain is a convergence region of distinct motor programs in larval locomotion.

## Results

### Building a template VNC coordinate for synapse registration

To map synapse configurations obtained from distinct immunostained samples into a common framework, a template coordinate system was generated. We labeled 60 VNCs with an antibody staining against cell adhesion protein Fasciclin2, which sparsely marks axon bundles and allows us to define reliable landmarks within the neuropil (13) (Figure 1A). We manually labeled 48 landmarks in each of the 60 VNC confocal images (Figure S1). These 3D images were matched by a rigid transformation (i.e., translation and rotation) to minimize the distance between the same landmarks in the 60 images. After the transformation, the coordinates of a landmark in all the images were averaged to obtain the coordinates of the landmark in the template VNC (Figure 1B; see Methods for detailed procedure.) Among the Fasciclin2-immunoreactive axon bundles, Transverse Projection 1 (TP1) sends dorsoventral projection at the midline between the posterior commissure at a neuromere and the anterior commissure at the next posterior neuromere (13). In this study, the location of TP1 was used as an anatomical segment boundary of the neuropil, as previously reported (13). Based on this definition, our template covered six abdominal neuromeres, A1 to A6 (Figure 1C). The template coordinates were used to register images of the following anatomical and activity data into the common coordinate space.

**Figure 1.**
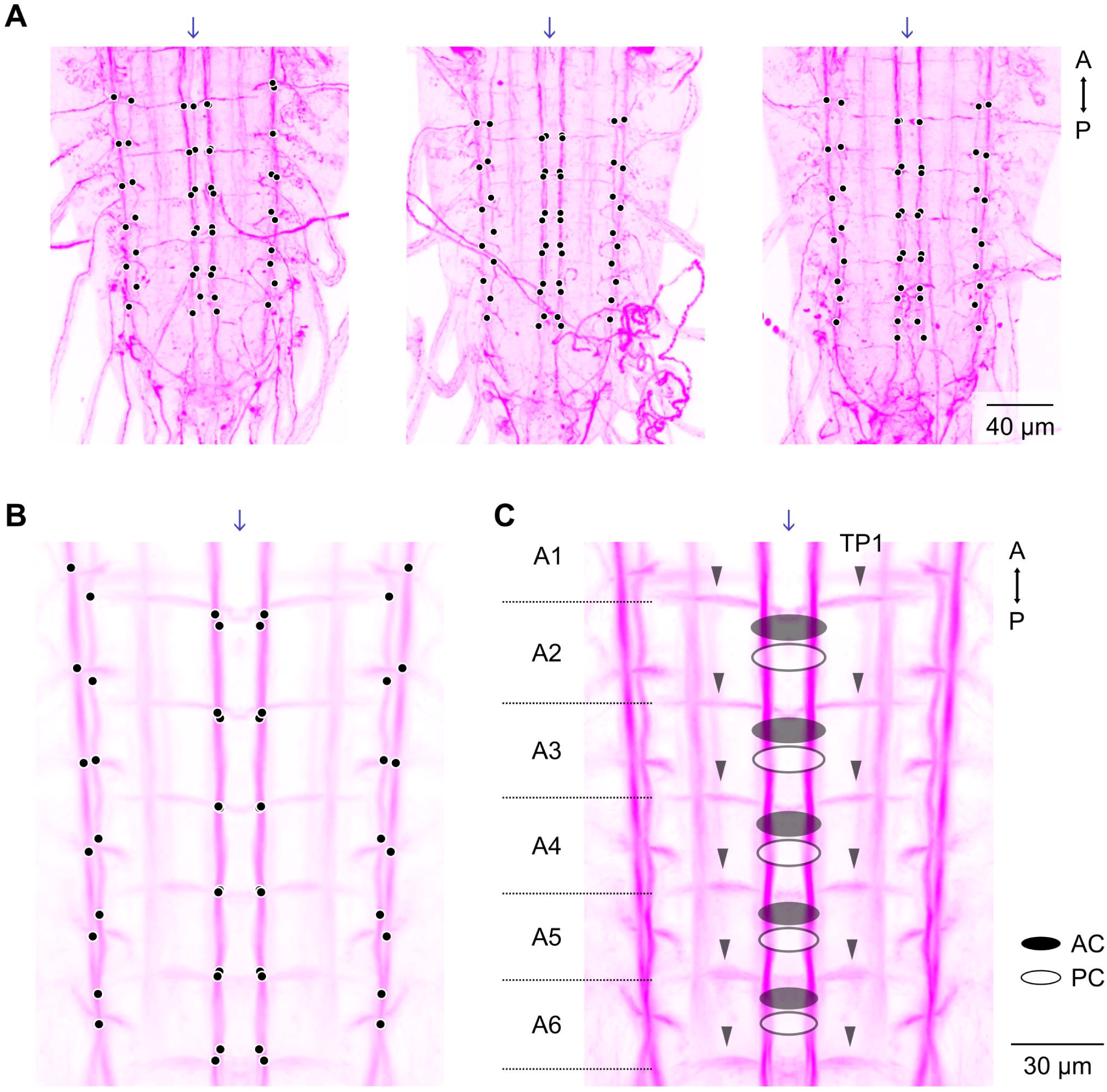
Construction of a spatial framework template based on Fasciclin 2 (Fas2) landmarks. (A) Three example samples of anti-Fas2 staining data. Black dots show 48 landmarks labeled manually for each sample. (B) Fas2 landmarks (black dots) in the template framework which were obtained by aligning and averaging the coordinates of the labeled landmarks in 60 samples. (C) The template framework covers six abdominal neuromeres (A1-A6). Black arrowheads indicate transverse projection 1 (TP1) of Fas2-positive bundles. AC: anterior commissure, PC: posterior commissure. Arrows indicate the midline.

### Expression patterns of neurotransmitter markers in the larval VNC

To analyze the spatial organization of synapses in the motor circuits, we examined the distribution of markers for fast-acting neurotransmitters. The fly nervous system uses three neurotransmitters for fast synaptic transmission: acetylcholine, glutamate, or gamma-aminobutyric acid (GABA) (14). We labelled these synapses with antibodies against neurotransmitter-specific proteins. Choline acetyltransferase (ChAT) encodes an enzyme to produce acetylcholine. Vesicular glutamate transporter (VGlut) and vesicular GABA transporter (VGat) encode transporters required for pumping glutamate and GABA into the lumen of synaptic vesicles, respectively (15,16). Confocal images of VNCs immunostained with each of these antibodies were registered into the template VNC (Figure 2A-2E). We noticed a difference in the use of neurotransmitters between the dorsal and ventral regions of the VNC. In the ventral region, the vast majority of synapses are GABAergic (Figure 2B-2C). In contrast, the three neurotransmitters are used in the dorsal region in similar amounts. Previous observations show that the mechanosensory and nociceptive neurons send projections to the ventral region, suggesting that the ventral region operates sensory processing (17,18). On the other hand, motor neuron dendrites and proprioceptive sensory axons target the dorsal region, which indicates that the dorsal region possesses motor control circuits (17,18). Our observation of the dorsoventral difference in the use of neurotransmitters suggests that the sensory processing is conducted mainly by GABAergic transmission, whereas the motor control circuits capitalize on the three fast-acting neurotransmitters.

**Figure 2.**
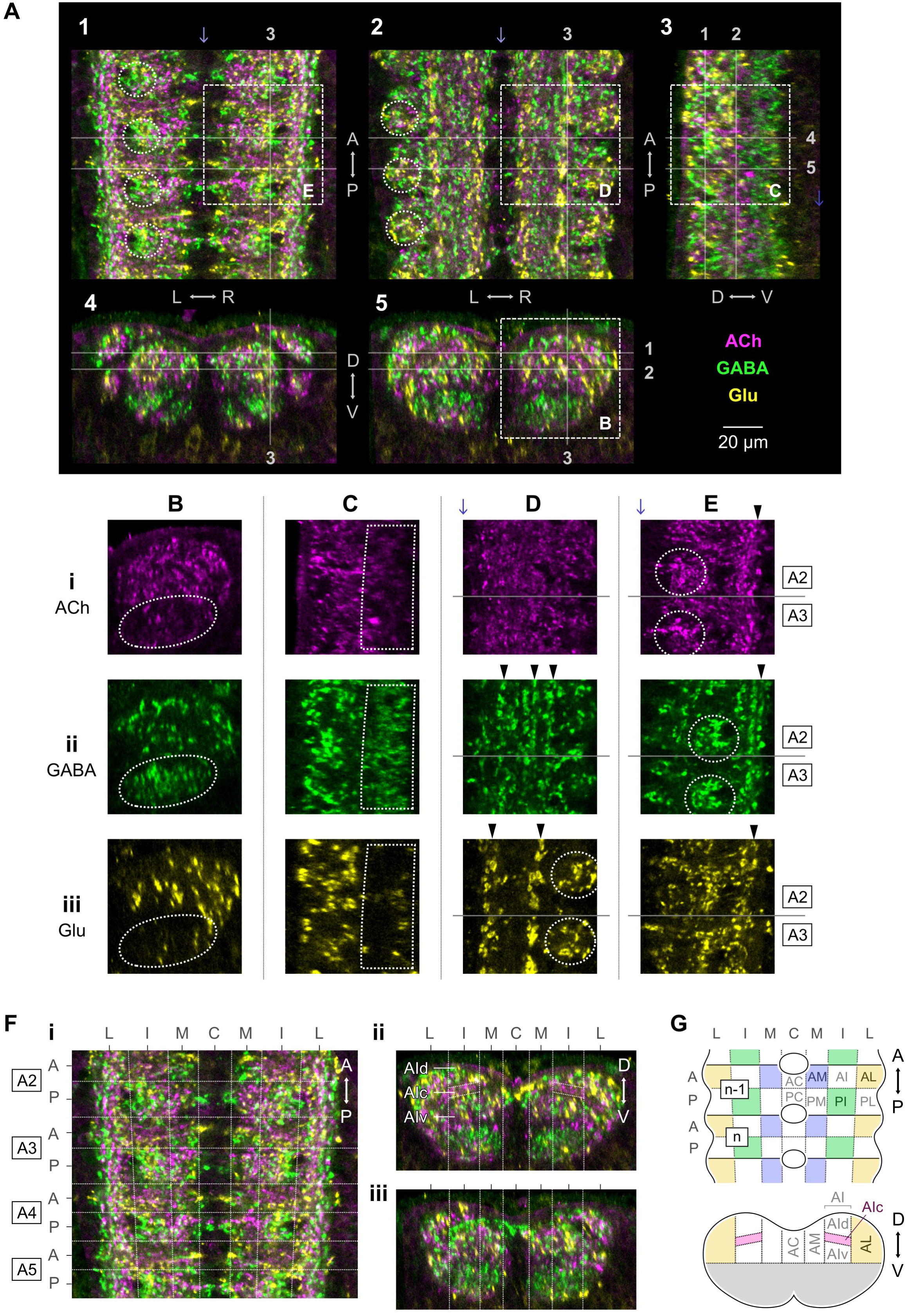
Distribution of fast-acting neurotransmitters in the neuropil. (A) Five views of a staining image in the template framework. Three distinct confocal images, anti-ChAT, anti-vGat, and anti-vGluT, are mapped onto the template coordinate and merged. 1, 2: dorsal sections, 3: sagittal section, 4, 5: frontal sections. Arrows indicate the midline. (B-E) Immunostaining images showing the areas indicated by squares in (A), with splitting the channels (i: ACh, ii: GABA, iii: Glu). (B, C) Dotted lines indicate GABA-rich ventral regions. (D, E) Black arrowheads indicate strips, while dotted circles indicate blobs. Blue arrows indicate the midline. (F) Spatial pattern of the neurotransmitters was subdivided into grid-wise areas. i: a dorsal section, ii: the frontal section centering the anterior half of the A4 neuromere, and iii: the frontal section centering the posterior half of the A4 neuromere. (G) Schematic diagram of 10 domains in the neuropil. Each hemi-neuromere is divided into anterior and posterior halves in the AP direction, and commissure, medial, intermediate, and lateral areas in the ML direction. Antero-intermediate regions are further subdivided into dorsal, central, and ventral areas in the DV direction.

To reveal synaptic organization in the motor circuits, we analyzed the distribution of neurotransmitters in the dorsal region of the VNC. We noticed two morphological patterns in the synapse distribution: strips and blobs (Figure 2D-2E). In the strips, synapses of the same neurotransmitters are stretched along the body axis in several locations. In the blobs, the same neurotransmitter markers are clustered with a diameter of half of the width of a neuromere. While the strips of distinct neurotransmitters are intermingled, the blobs of distinct neurotransmitters are located at different regions in the VNC (See circles in Figure 2A, 2D and 2E). This observation implies that the scale of these blobs could be a unit for motor control (Figure 2F). In addition, in the frontal view, we noticed that the anterior intermediate part of a hemi-neuromere includes a stripe where the cholinergic marker dominates (Figure 2Fii). To analyze the structural and functional significance of these partitions, we demarcated a hemi-neuromere into ten intrasegmental domains (AC, AM, AId, AIc, AIv, AL, PC, PM, PI and PL) (Figure 2G) and analyzed the properties of these domains below.

### Correspondence between the intrasegmental domains and the myotopic map

First, to reveal the domains involved in the motor output, we mapped the dendrites of motor neurons onto the template VNC based on connectomics data established from transmission electron microscopic images of a serial sectioned larval CNS (19,20). We manually located the landmarks for the VNC registration in the electron microscope images and mapped the skeleton data of neurons by non-linear transformation (See Methods section for detailed procedure). It should be noted that while we used the third instar larvae in this study, the connectomics data was generated from a first-instar larva. Although the size of the CNS in the third instar is larger than that in the first instar, the connectivity topography in a circuit is conserved across larval development (21). Accordingly, the connectomics data of neurons in the first instar was used to study the domains defined in the third instar.

The dendrites of larval motorneurons form the myotopic map, where the location of motoneuronal dendrites is partitioned based on the target muscles (18). This myotopic map is observed in the template VNC (Figure 3). Three classes of motor neurons (ISN, ISNb, and ISNd motor neurons), which innervate longitudinal muscles (13,22), form dendrites centered around the posterior-intermediate (PI) region. The other two motor neuron classes (SNa and SNc motor neurons), which target transverse muscles (13,22), extend the dendrites around the anterior-lateral (AL) region. This observation implies that the scale of the intrasegmental domain based on the neurotransmitter expression (Figure 2F) could serve as a unit for motor control. The layout of motor neurons innervating a single body wall segment is parasegmental: the body wall muscles in a segment are targeted either by SNa and SNc motor neurons in the same VNC segment or by ISN, ISNb, and ISNd motor neurons in the segment next anterior (18). This property is observed in the template VNC data (Figure 3C-3D). These observations suggest that four intrasegmental domains in the n-1th segment (PC_n-1, PM_n-1, PI_n-1, and PL_n-1) and six domains in the nth segment (AC_n, AM_n, AId_n, AIc_n, AIv_n and AL_n) could be grouped as a segmental unit for controlling body wall muscles in a single hemi-segment. Furthermore, connectomics data suggest that the PI and AL domains comprise most of the dendrites of motor neurons (Figure 3E).

**Figure 3.**
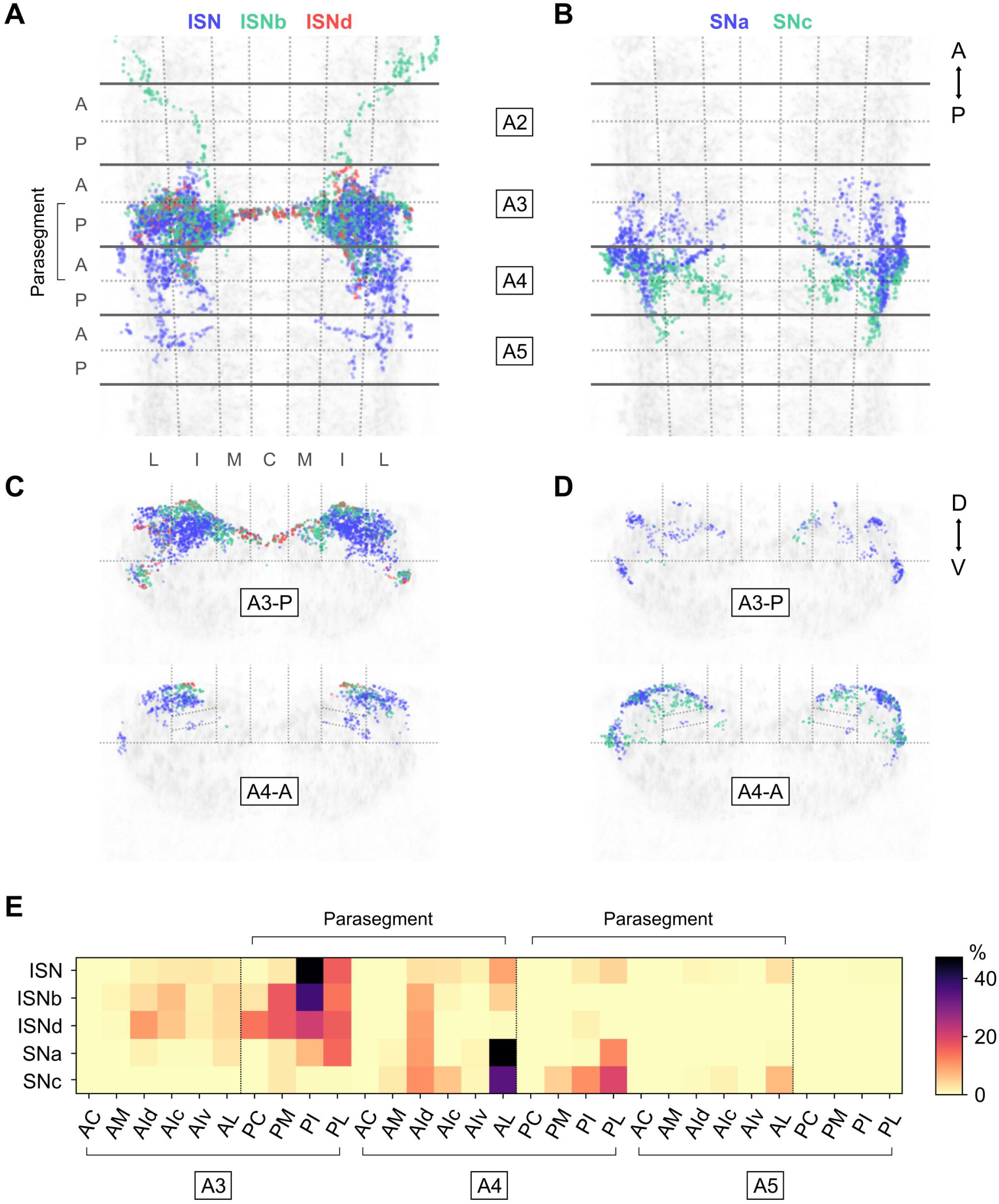
Relationship between postsynaptic terminals of motor neurons from connectomics data and the intrasegmental domains. (A and B) Postsynaptic terminals of ISN, ISNb, and ISNd motor neurons (A) or SNa and SNc motor neurons (B) are mapped onto the template framework in the dorsal view. Solid and dotted lines indicate neurotransmitter domains defined in Figure 2F. (C and D) Postsynaptic terminals of ISN, ISNb, and ISNd motor neurons (C) or SNa and SNc (D) in the posterior half of the A3 neuromere (top) and the anterior half of A4 neuromere (bottom) are shown in frontal view. (E) Distribution of postsynaptic sites in the neurotransmitter domains for each motor neuron group. Percentage indicates the number of postsynaptic terminals in a domain normalized by the total for each motor neuron group.

### One domain pair shows a fixed order in activity recruitment in both forward and backward waves

Next, to examine the functional significance of the intrasegmental domains, we analyzed synaptic activity within the segmental units defined above (Figure 3E). To this aim, we conducted calcium imaging of the isolated VNC, which shows fictive locomotion (23,24). We expressed a membrane-bound form calcium indicator in all neurons, recorded fluorescence signals with an EMCCD camera, and extracted calcium signals from bouton-like structures (25). After the calcium imaging, the samples were fixed and immunostained with the antibody against Fasciclin2 and an antibody against GFP and scanned by a confocal microscope. The calcium imaging data were mapped into the template VNC coordinate with manually labeled landmarks (See “Spatial matching of calcium imaging to immunostaining data (GFP matching)” in Methods for detailed procedure) (Figure S2). Corresponding to forward and backward crawling, the isolated larval VNC exhibits the propagation of synaptic activity from posterior to anterior and from anterior to posterior, respectively.

We compared the activity timings of the domains between forward and backward waves. In forward waves, the activities among domains are not synchronized but have a delay (Figure 4A). The similar tendency is observed in backward waves, but the order of activity recruitment depends on the pairs (Figure 4B-4C). To analyze the recruitment order between domains, we plotted the delays in forward and backward waves for each pair (Figure 4D). The plot shows that while most pairs exhibit the opposite order between forward and backward waves, two pairs of domains within the segmental unit show the same recruitment order: PI_n-1 and PL_n-1 are followed by AIc_n in both forward and backward waves. Especially, the delay from PI_n-1 to AIc_n does not significantly differ between forward and backward, unlike PL_n-1 to AIc_n, which indicates that the activity propagation of PI_n-1 and AIc_n is consistent between the two motor patterns (Figure 4E; AIc_n – PI_n-1: 0.35 ± 0.43 s for forward, 0.55 ± 0.31 s for backward (mean ± std.)). As shown above (Figure 3C), the PI domain is the primary region that generates output for motor neurons that innervate longitudinal muscles. These observations suggest that the PI domain, which is involved in driving forces for propulsion, and the AIc domain show the fixed sequence of activity in the waves in both directions.

**Figure 4.**
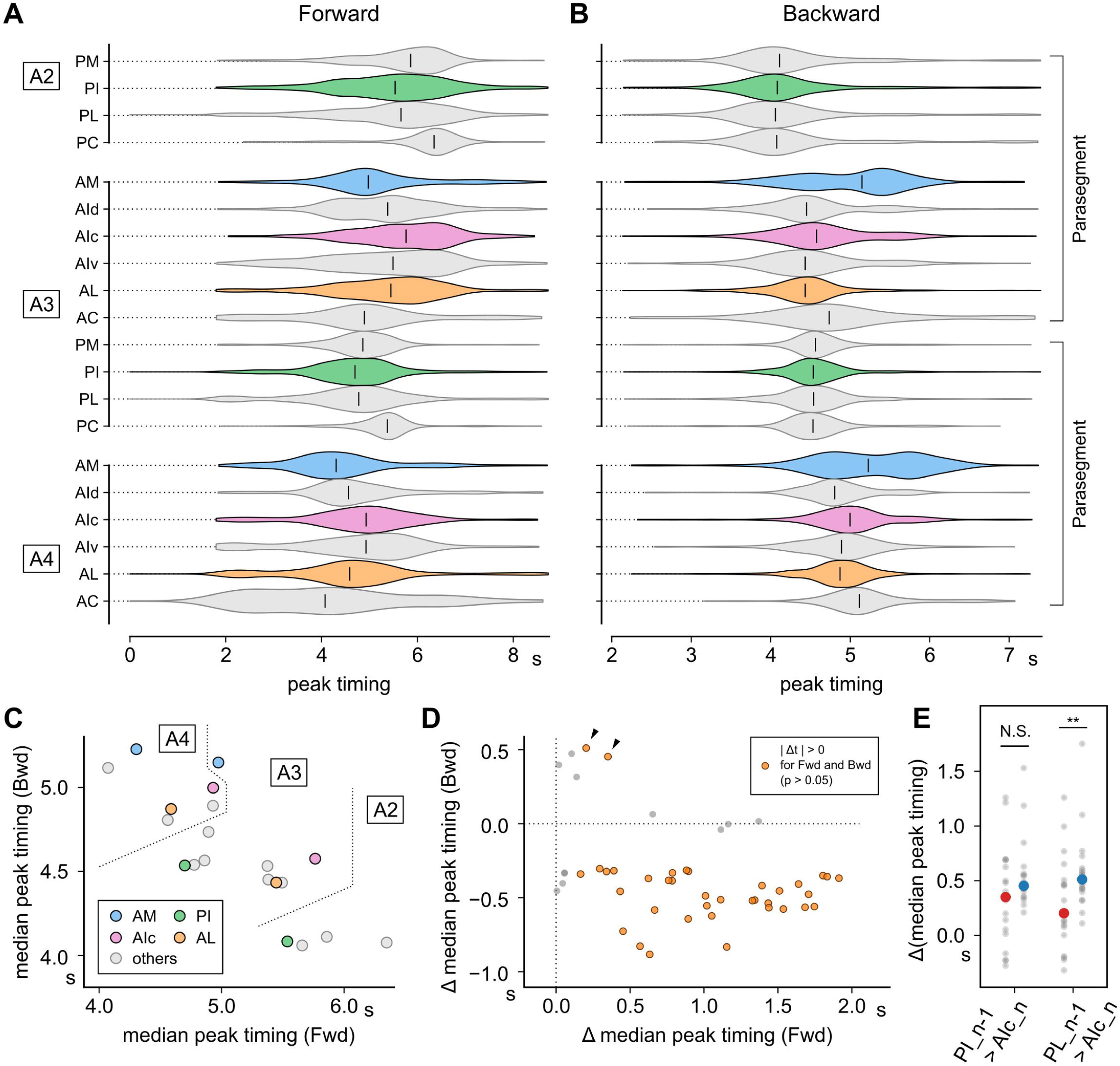
PI and AIc domains show consistent activity delay in forward and backward waves. (A and B) Activity peak timings of extracted boutons in each domain during an event-averaged forward wave (A) or backward wave (B). (C) Scatter plot showing median peak timings of domains (A) and (B). Horizontal and vertical axes indicate median peak timings during forward and backward waves, respectively. (D) Scatter plot showing time lags (difference of median peak timings) for every pair of domains in the parasegmental section. Horizontal and vertical axes indicate time lags during forward and backward waves, respectively. Two pairs of domains explicit activity lag in the same direction for forward and backward waves (black arrowheads). (E) Activity delays of the two domain pairs indicated by black arrowheads in (D) are compared. Gray dots indicate time lags of median peak timing in domain pairs (PI_n-1 to AIc_n, or PL_n-1 to AIc_n) for every ipsilateral combination from every sample. Red and blue dots are the median of gray dots. **: p < 0.01 (Wilcoxon rank sum test).

### Interneurons that form presynaptic terminals in the PI and AIc domains

To identify interneurons involved in the activity of the PI and AIc domains, we searched for interneurons that have pre-and post-synaptic terminals in these domains (Figure 5). The PI domain consists of the presynaptic terminals of A23a and A31k GABAergic neurons (26) (Figure 5A). These neurons should be the origin of GABAergic blobs observed in immunostaining (Figure 2F). The immunostaining data show that the PI domain also possesses cholinergic and glutamatergic neurons (Figure S3). Consistent with this, the connectomics data indicate that A03 (20), A08 (27), and A18 lineages (28) include cholinergic neurons targeting the PI domain, and glutamatergic neurons targeting the PI domain includes A02e (26,29) (Figure 5B). Although the origin of ChAT positive signal in the AIc domain remains unclear, the AIc domain includes postsynaptic terminals of cholinergic neurons A01c and A01ci (26) (Figure 5C). This observation suggests that the PI and AIc domains form functional units for controlling larval crawling behavior.

**Figure 5.**
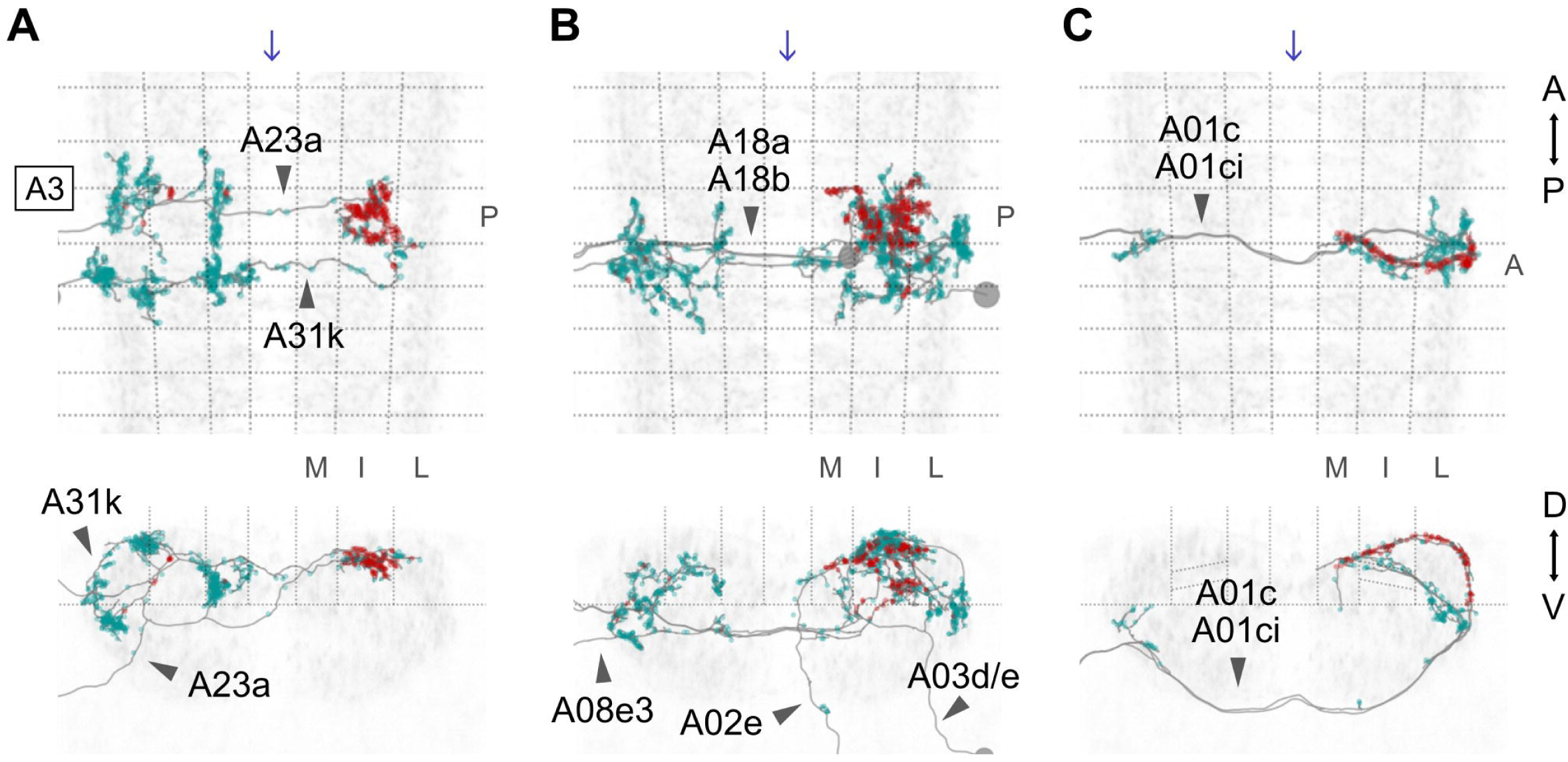
Pre- and post-synaptic terminals of interneurons involved in motor coordination are consistently distributed with the neurotransmitter domains. Interneurons reconstructed from the connectomics data are mapped onto the template framework. Red and cyan dots indicate pre- and post-synaptic terminals, respectively. (A) A23a and A31k neurons. (B) A03d/e, A08e3, A18a, A18b, and A02e neurons. (C) A01c and A01ci neurons. Arrows indicate the midline.

## Discussion

### Synapse organization for multi-functional larval locomotion

In this study, we analyzed the anatomical and functional organization of synapses for fly larval locomotion. The motor circuits consist of the three fast-acting neurotransmitters in similar amounts in contrast to GABA-dominant sensory circuits. The hemi-neuromere of the motor circuit was partitioned into ten domains based on the distribution of the neurotransmitter markers. Calcium imaging analysis shows that one pair of the ten domains, the posterior-intermediate domain at the nth segment (PI_n) and the central anterior-intermediate domain at the n-1th segment (AIc_n-1), have a consistent order in the activity recruitment in forward and backward waves whereas the other pairs are not.

The property of the recruitment order in PI_n and AIc_n-1 activity being independent of wave direction (forward vs backward) may underlie the multi-functionality of larval locomotion. There are about thirty body wall muscles in a hemisegment (18,22), and the sequence of contraction of individual muscles within a segment is similar between forward and backward (9) while some muscles show distinct contraction timing depending on the crawling direction (11). Accordingly, there should be direction-independent circuits to generate forward and backward crawling. Consistent with this notion, the PI_n domain contains presynaptic terminals of interneurons that have been reported as being involved in motor control (Figure 5A-5B). In contrast, we could not find domains recruited earlier than PI_n consistently in both forward and backward waves. These observations imply that PI_n is a convergence domain of forward and backward-engaging circuits, and the two distinct behaviors, forward and backward crawling, capitalize on the shared circuits formed in PI_n. Further analysis of upstream neurons to PI_n will provide a novel insight into how the distinct motor programs converge in neural circuits.

### The function of the domains exhibiting direction-independent recruitment activity

One of the direction-independent domains, the PI domain, has a blob of GABAergic terminals (Figure 2F) and presynapses of Glu and ACh (Figure S3). The existence of GABAergic clusters in the PI domain, where motor neuronal dendrites accumulate, implies two non-exclusive possibilities: (1) The GABAergic terminals are involved in the cooperative suppression of motor output. Blocking excessive contraction is critical for larval motor control (30). Coordinated activation of the GABAergic terminals may be involved in the well-timed motor suppression. (2) The GABAergic terminals trigger the excitation of motor neurons by post-inhibitory rebound. Larval motor neurons have the property to show excitation after hyperpolarization (31). Coordinated activity of the three fast-acting neurotransmitters would drive the contraction and relaxation of longitudinal muscles.

The other domain, AIc, contains the postsynaptic terminals of A01c and A01ci. These cholinergic neurons are upstream of transverse muscles (26). Accordingly, the AIc domain, which is recruited after PI in the segment next anterior, may be involved in the contraction of transverse muscles. Since transverse muscles contract after longitudinal muscles in an experimental condition (9), the direction-independent sequential activation of PI_n-1 to AIc_n would be involved in intrasegmental coordination of muscle contraction. In contrast, anatomical analysis shows the dendrites of motor neurons for transverse muscles (SNa and SNc) locate mainly in the AL domains (Figure 3D-3E). Calcium imaging assay demonstrates that the AL domains do not exhibit direction-independent activity recruitment with the PI domains (Figure 4A-4B). Since the contraction of transverse muscles during crawling depends on experimental conditions (32), the function of AIc activity is still unclear. Further connectomics and functional analysis of the PI and AIc domains will clarify the roles of direction-independent domains in motor control.

## Methods

### Drosophila melanogaster strains

All animals were raised on standard cornmeal-based food. For pan-neuronal calcium imaging, we used a fruit fly line of genotype *UAS-CD4::GCaMP6f* (26); nSyb-Gal4 (nSyb-Gal4: Bloomington #58763). For targeting specific neurons, we used the following drivers. Gal4 drivers (33,34): NP6051-Gal4 (Kyoto stock center #NP6051), R36G02-Gal410 (Bloomington #49939), and R75H04-Gal4 (Bloomington #39909).

### Calcium imaging

The CNS of third-instar fruit fly larvae was isolated by dissecting the animals with microscissors (29) was mounted on an adhesive slide glass (MAS-coated slide glass S9215, Matsunami Glass, Japan) and soaked in an insect saline (TES buffer: TES 5 mM, NaCL 135 mM, KCl 5 mM, CaCl2 2 mM, MgCl_2_ 4 mM, sucrose 36 mM; pH = 7.15). Fluorescence signals from the specimens were recorded by an EMCCD camera (iXon+ DU-897E-CS0-#BV, Andor, UK; 63x, 0.27 μm/px, exposure = 30 ms, EM gain = 144, data depth = 14 bit) with attached spinning disk confocal unit (CSU21, YOKOGAWA, Japan). The specimens were illuminated by a blue laser (CSU-LS2WF, Solution Systems, Japan; power = 300 to 500 μW, wavelength = 488 nm) through a water immersion objective lens (ACROPLAN 63X, Zeiss, Germany). During recordings, we scanned the dorsal-half VNC neuropil by quick vibration of the objective lens. The vibration was controlled by a piezoelectric device (P-725.2CL, PI, Germany). For each recording, five parallel focal planes with 5 μm intervals were alternately focused, where frames were snapped by 3 Hz for each plane.

### Bouton extraction from calcium imaging data

To analyze population dynamics in the neuropil, we extracted bouton-like regions in calcium imaging data. For bouton extraction, we designed an algorithm to decompose neuropil imaging data into bouton-sized pixel clusters (PQ-based clustering (25)). In brief, the algorithm searches the optimal configuration of pixel clusters in which pixels show relatively similar activity traces. For this purpose, the similarity network among pixels is used to calculate an expanded version of modularity (35), which is defined in graph theory. Modularity is a function of clustering configuration, and the better clustering configuration is supposed to provide a larger quantity of modularity. Modularity is maximized by the simulated annealing method, in which the temperature of the system is gradually reduced during thermal fluctuation of system states depending on energy function as modularity × (- 1).

### Immunohistochemistry

Immunohistochemistry was performed on isolated CNSs of third-instar larvae. Samples were fixed with 3.7 % formaldehyde for 30 min at room temperature, washed with 0.2 % Triton X-100 in PBS for 30 min at room temperature, and blocked with normal goat serum for 30 min at room temperature. The processed samples were incubated at 4 °C for at least two days in each primary and secondary antibody solution. Primary antibodies: rabbit anti-GFP (Frontier science #Af2020, 1:1000), rabbit anti-HA (Cell Signaling Technology #C29F4, 1:1000), rabbit anti-VGAT (16) (1:300), rabbit anti-VGluT (15) (1:1000), mouse anti-ChAT (DSHB #4B1, 1:50), and mouse anti-Fas2 (36) (DSHB #1D4, 1:300). Secondary antibodies: Alexa-Fluor 488-conjugated goat anti-rabbit IgG (A-11034, 1:300) and Alexa-Fluor 555-conjugated goat anti-mouse IgG (A-21424, 1:300). The stained samples were recorded with a confocal microscopy (BX61WI + FluoView FW1000, Olympus, Japan) and an oil immersion objective lens (Plan-APOCHROMAT 63X, Zeiss, Germany).

### Spatial matching

Spatial matching was performed between multiple types of datasets recording the VNC neuropil. We started the process with manual labeling of reference points at the same locations in the query and target dataset and then generated coordinate transforms in which every labeled point in the query dataset was mapped onto each corresponding point in the target, where the transforms were provided by thin-plate spline (TPS) method (37,38). Since TPS defines smooth and natural one-to-one mapping between the query and target coordinate systems, any locations in the query data could be assigned to coordinates in the target. TPS was calculated by an open-source Python library (https://github.com/tzing/tps-deformation). To enhance the preciseness of point labeling, we designed a GUI platform for viewing arbitral sections of three-dimensional images. The platform provided an intuitive user interface to manipulate the position of a plane to display a section and label points on the section with three-dimensional coordinates. The software is released online with a brief installation guideline (https://github.com/Fukumasu/SectionViewer).

### Spatial matching of calcium imaging to immunostaining data (GFP matching)

Calcium imaging data and immunostaining data of the same sample were spatially matched to precisely identify the location of calcium imaging frames in the VNC neuropil. Since the data were acquired from identical samples, it was possible to locate multiple pairs (15–40 pairs per sample) of corresponding points between imaging and staining data by manually comparing those spatial patterns of GCaMP and anti-GFP fluorescence signals, respectively. To test the accuracy of the transforms calculated by TPS with the reference points, we applied leave-one-out cross-validation to the point sets. The mean error distance was 1.8 ± 1.2 (mean ± std., n = 121 from five samples) µm, which indicated that the accuracy of transforms reached the level of bouton size (∼2 µm).

### Spatial matching of immunostaining to template coordinate system (Fas2 matching)

To align the immunostaining data of distinct samples, a template coordinate system was created, and the samples were spatially matched to the template. To label common points among the samples, we referred to the configuration of Fas2 bundles in the neuropil. We labeled reference points on longitudinal projections of the Fas2 bundles (DL, VL, DM and VM). For DL and VL, reference points were taken as intersections of these longitudinal bundles and planes defined by left and right TP2. For DM and VM, intersections of these bundles and planes defined by left and right TP1 and TP3 were used as reference points. The reference points were taken from six segments (A1–6) and left and right hemi-segments. In total, 4 (DL, VL, DM and VM) × 6 (A1–6) × 2 (left and right) = 48 points were labeled in each sample. By processing the coordinates of reference points from 60 samples, we defined a template set of 48 reference points. To make the template set of points symmetric according to the median plane, we generated an additional 60 sets of points by duplicating and mirroring the original 60 samples. The 120 sets of points in total were moved and rotated in three-dimensional ways to overlap with each other as closely as possible, and those coordinates were averaged. Since it was difficult to find the optimal positions of the point sets at once, the optimization was conducted iteratively. First, the mean coordinates (x, y, and z) of the point sets were calculated (mean point set). Then, each point set was rotated to match the mean point set as closely as possible, and mean coordinates were calculated again from the updated point sets. By repeating these steps, the point sets gradually approached a common configuration. After three iterations, the coordinate variation of the point sets was sufficiently converged, and we defined the final mean point set as the template. Transforms from immunostaining samples to the template coordinate system were calculated by TPS between reference points of the samples and the template. Leave-one-out cross-validation was applied to test the accuracy, and the error was 1.7 ± 1.1 (mean ± std., n = 2,880) µm, indicating that the matching was sufficiently accurate to compare samples in the resolution of bouton-size (∼2 µm).

### Spatial matching of ssTEM dataset to template coordinate system

Reconstructed skeletons of neurons in the ssTEM dataset were spatially matched to the template coordinate system. Acquisition and analysis of the ssTEM data have been reported (19,20,27,39). Whereas calcium imaging in the current study revealed the spatiotemporal structure of the A3 segment in third-instar larvae most clearly, neuronal reconstruction in the ssTEM data of a first-instar larva has been concentrated on the A1 segment. To spatially align segments from these frameworks in the best combination, we matched the A1 segment in the ssTEM data and the A3 segment in the template coordinate system based on the observation that the configuration of neurons in the neuropil is robustly conserved among distinct stages of larvae and also among segments A1–7. For reference points to calculate TPS, we manually labeled corresponding 38 locations on skeletons in the ssTEM data (A01ci, A02e, A23a, A27h, A27k, and A31k) in the A1 segment and the same neurons in the A3 segment mapped onto the template coordinate system by Fas2 matching. The mean error of the matching calculated by leave-one-out cross-validation was 4.0 ± 1.8 (mean ± std., n = 38) µm, which indicated sufficient accuracy to compare branches in reconstructed skeletons and the intra-segmental domains.

### Temporal matching of bouton activities

Temporal matching was applied to activity profiles of boutons to temporally merge distinct events and samples. Temporal matching was first performed among FW or BW events of the same sample, and then the averaged events of multiple samples were temporally aligned. For both cases, the speed of wave propagation was normalized based on linear regression of recruitment timing and AP position in the template space of boutons, and new activity traces were generated by interpolation based on the discrete Fourier transform. Recruitment timing was defined as the time point of the maximum slope in each activity trace (40). Timing of the maximum slope was estimated as the inflection point of the cubic function passing through four data points around the maximum slope.

## Acknowledgment

We thank Bloomington Drosophila Stock Center and KYOTO Drosophila Stock Center for the fly lines. We thank Developmental Studies Hybridoma Bank, Dr. Hermann Aberle and Dr. David Krantz for the antibodies. We thank Dr. Albert Cardona for continued access to the L1 EM dataset. This work was supported by MEXT/JSPS KAKENHI grants (17K19439, 19H04742, 20H05048, 21H02576 21H05675, 22K19479, 22H05487, 23H04213, 24H01225, 24K02117 to A.N. and 17K07042, 20K06908, 21H05301 and 23K05959 to H.K.).

## Competing interests

We have no conflicts of interest with respect to the work.

## Notes

### Competing Interest Statement

The authors have declared no competing interest.

### Summary of Updates

The Acknowledgements section has been revised.

