## Supplementary figures and images for "Modular organization of synapses within a neuromere for distinct axial locomotion in *Drosophila* larvae"

### Figure S1

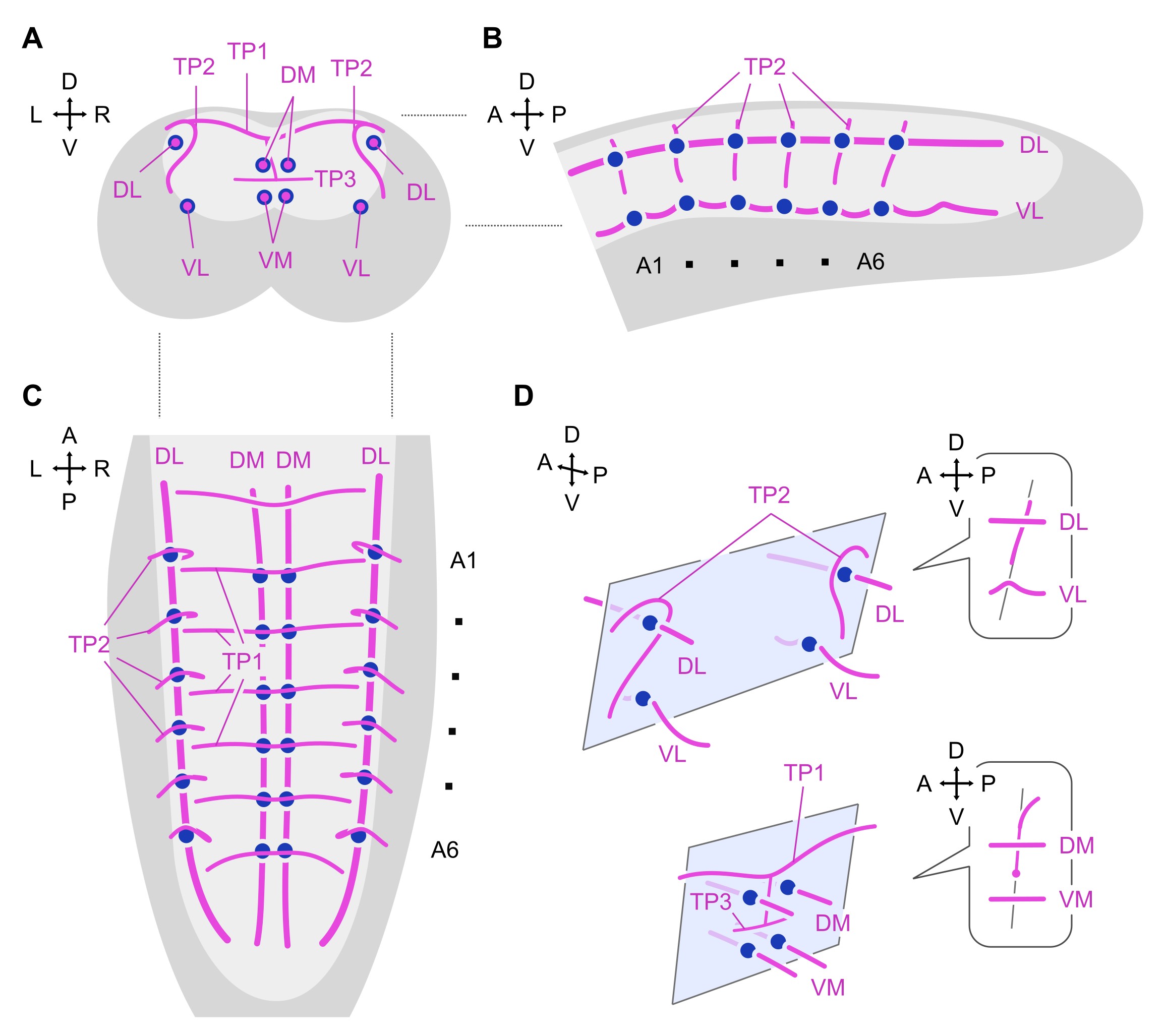

### Figure S2

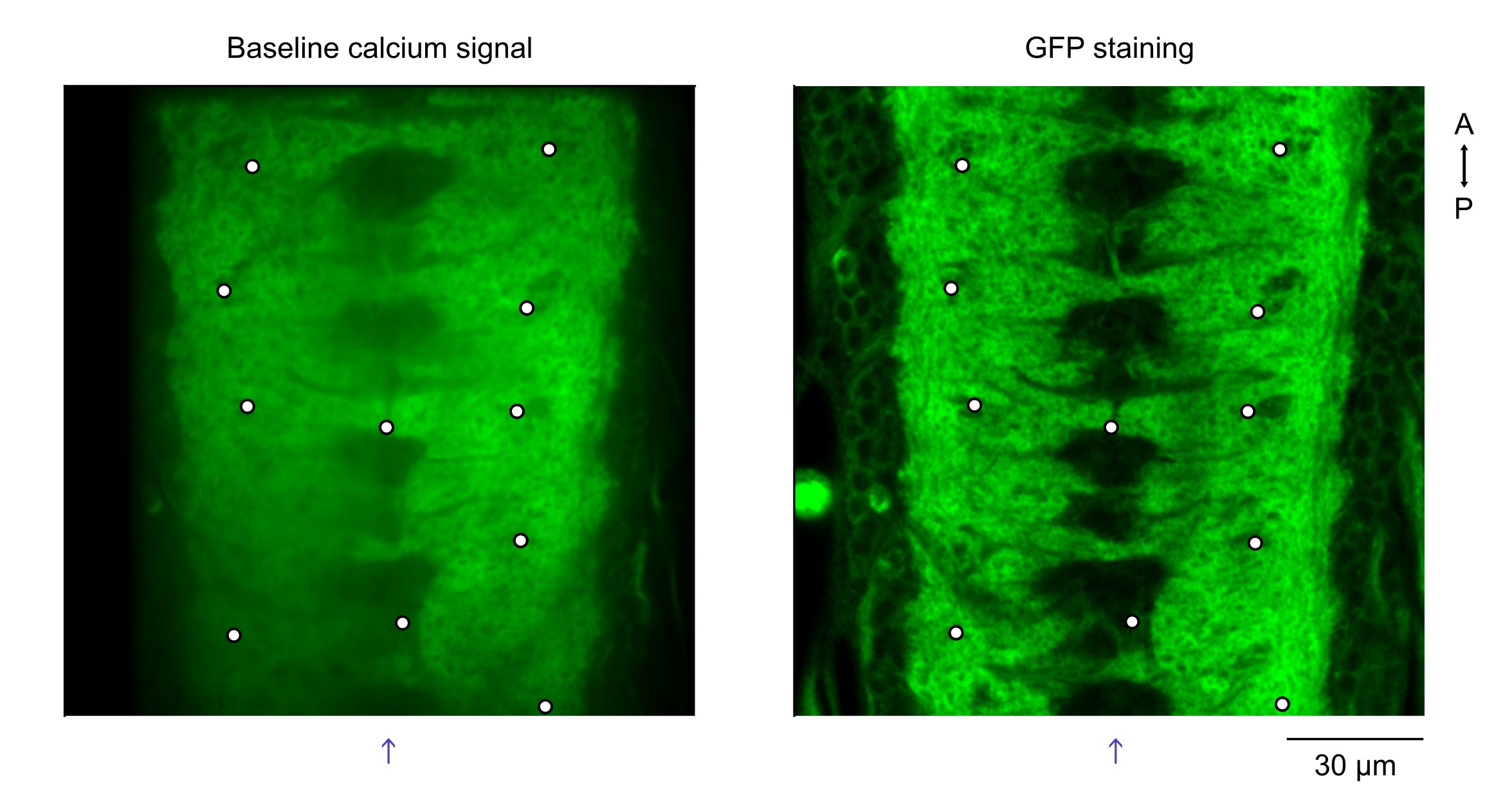

### Figure S3

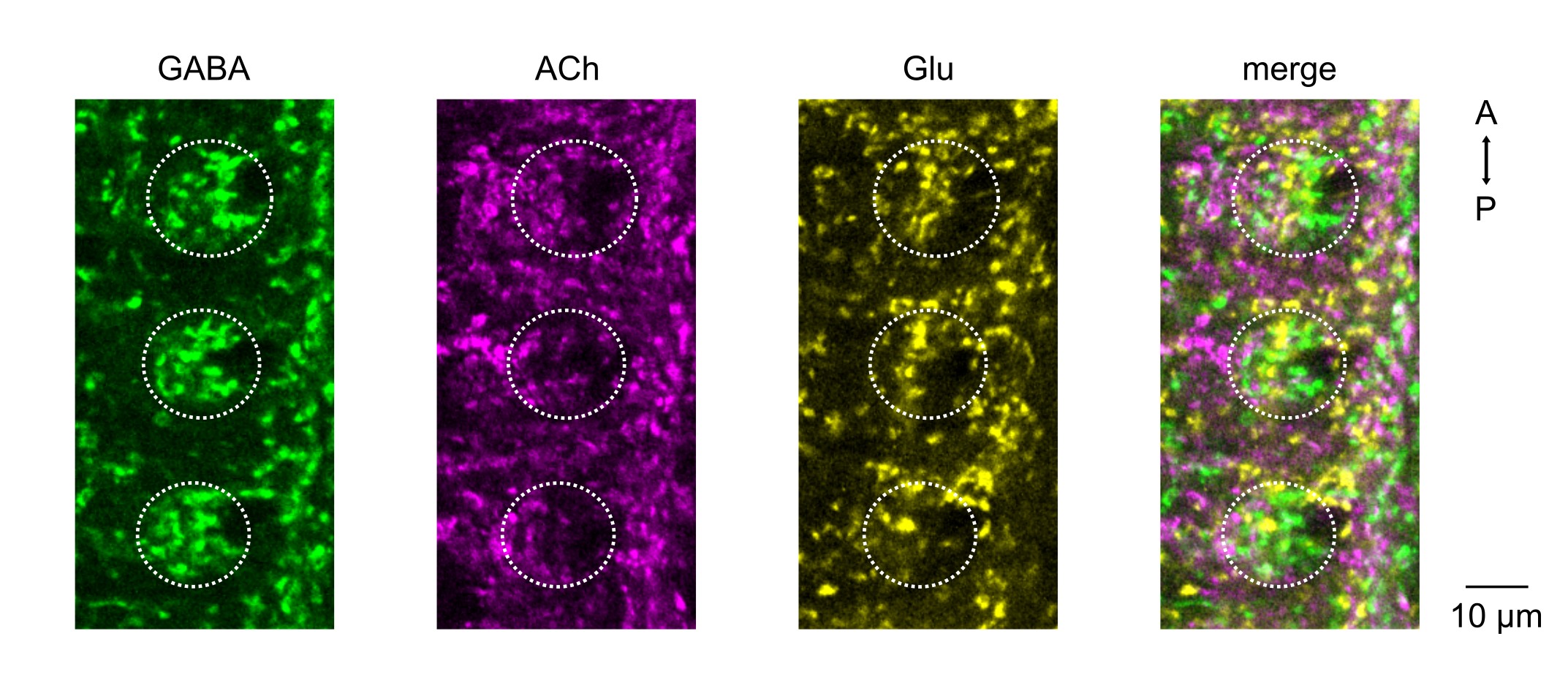
